## Supplemental Figures for "A scheduler for rhythmic gene expression"

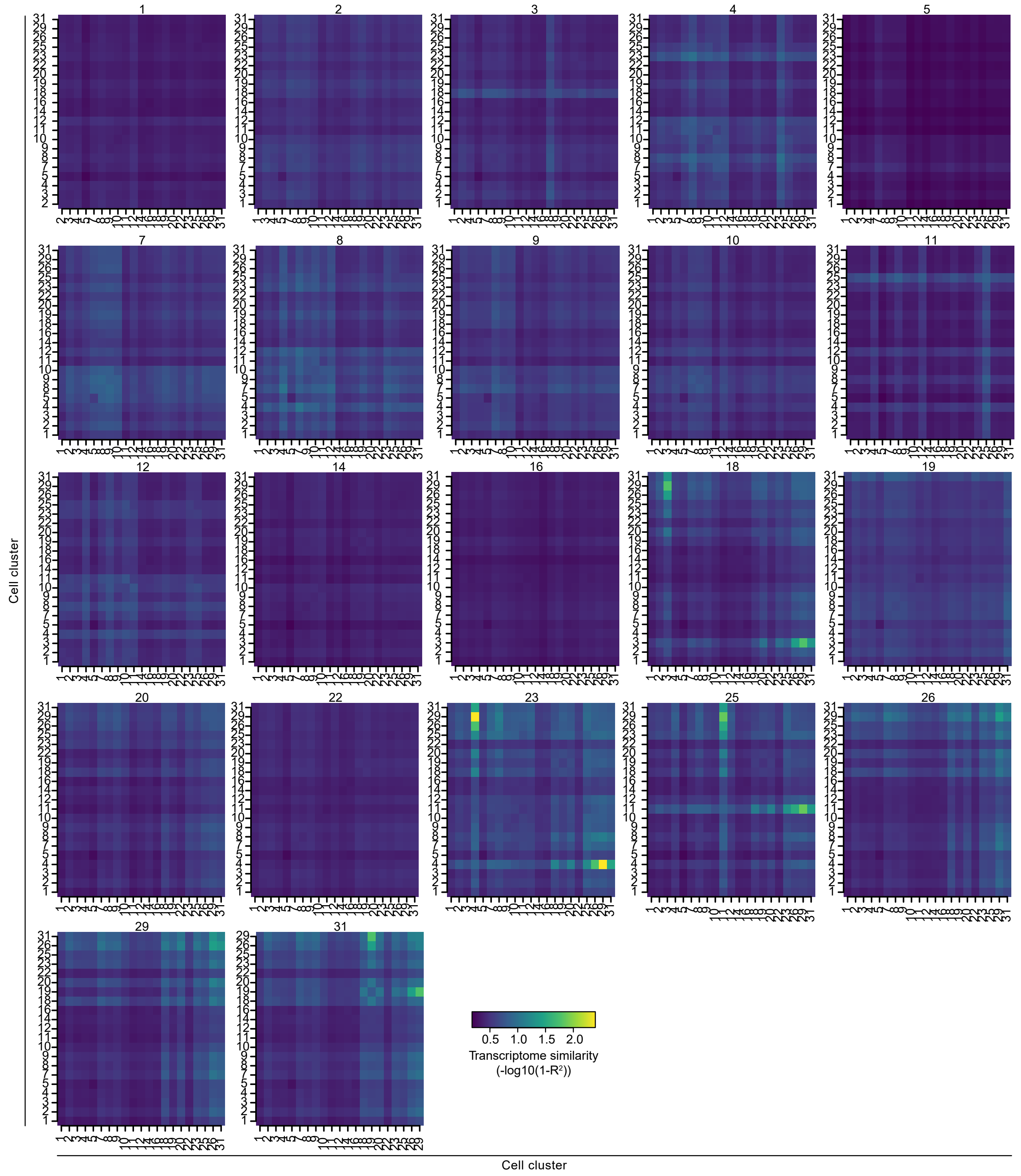


**Supplementary Figure 1**

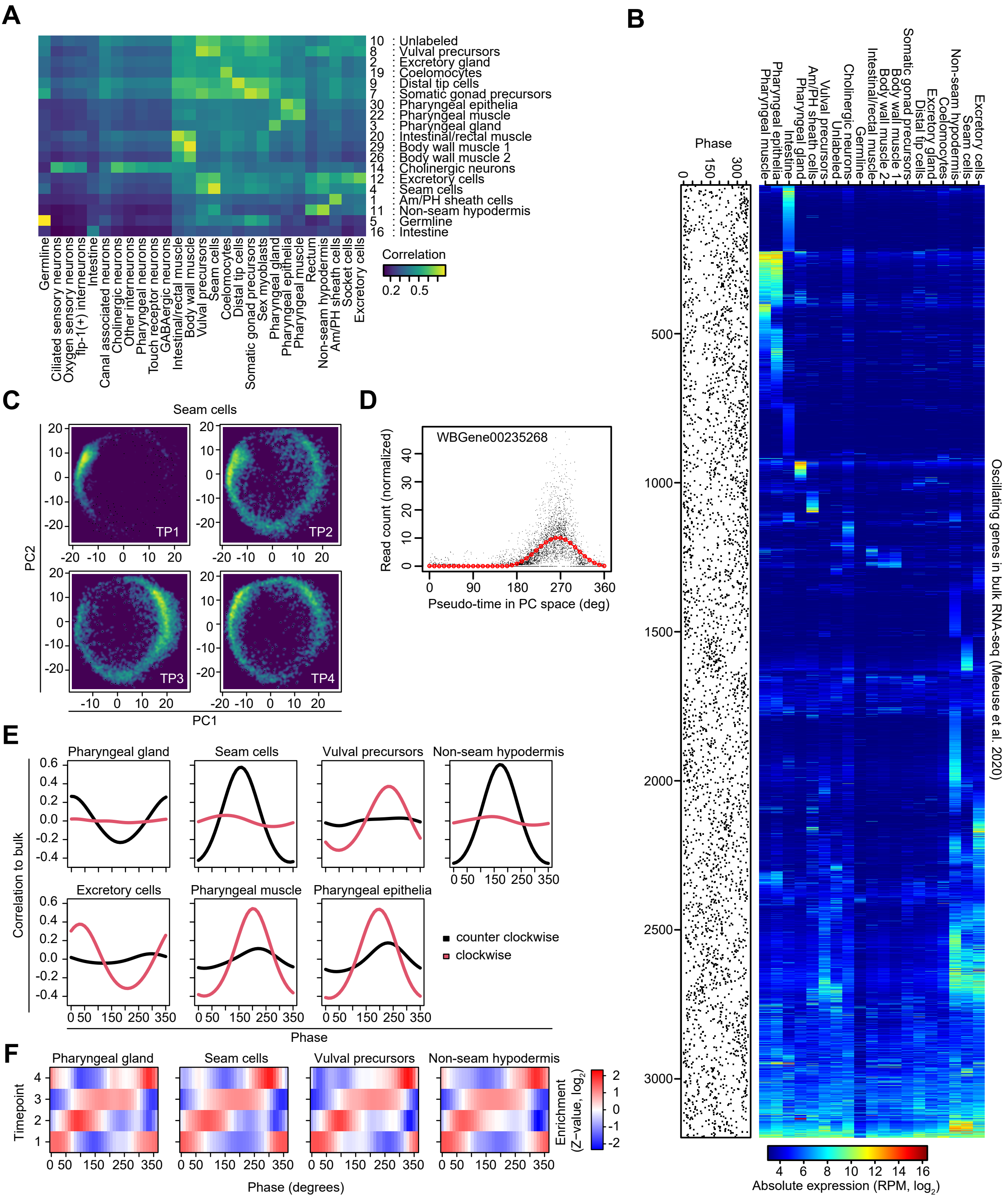


### Supplementary Figure 2

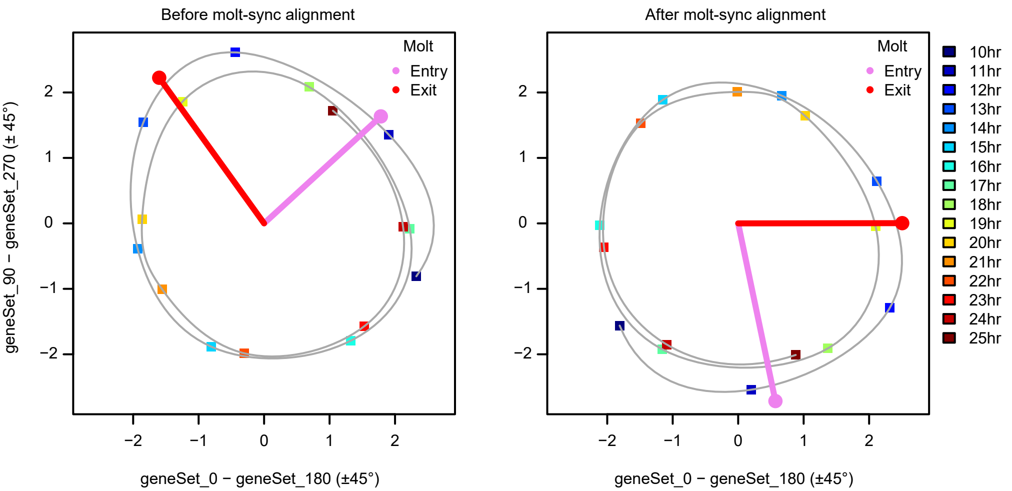


**Supplementary Figure 3**

**A**

| Tissue | #Cells |
|---|---|
| Am/PH sheath cells | 4 |
| Excretory gland | 2 |
| Pharyngeal gland | 5 |
| Seam cells | 32 |
| Germline | N/A |
| Somatic gonad precursors | 10 |
| Vulval precursors | 6 |
| Distal tip cells | 2 |
| Unlabeled | N/A |
| Non-seam hypodermis | 165 |
| Excretory cells | 3 |
| Cholinergic neurons | N/A |
| Intestine | 32 |
| Coelomocytes | 6 |
| Intestinal/rectal muscle | 4 |
| Pharyngeal muscle | 37 |
| Body wall muscle 2 | N/A |
| Body wall muscle 1 | 95 |
| Pharyngeal epithelia | 9 |

**B**

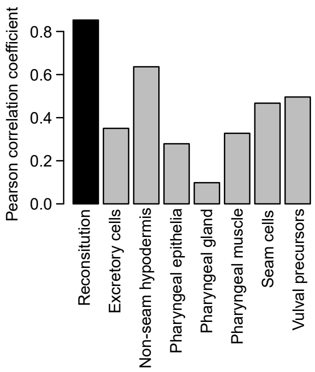


**C**

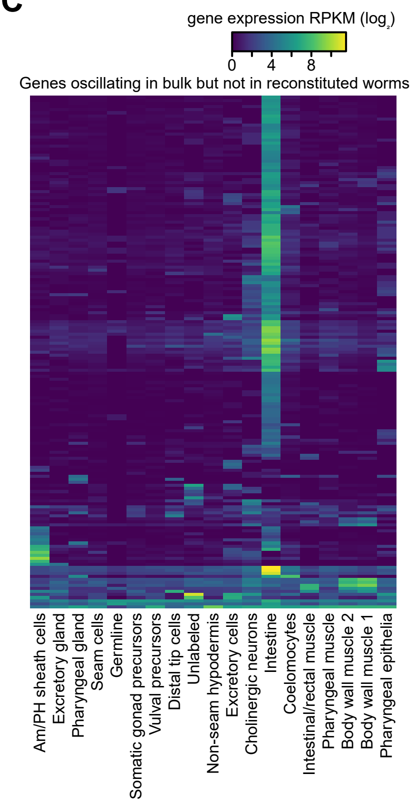


**Supplementary Figure 4**

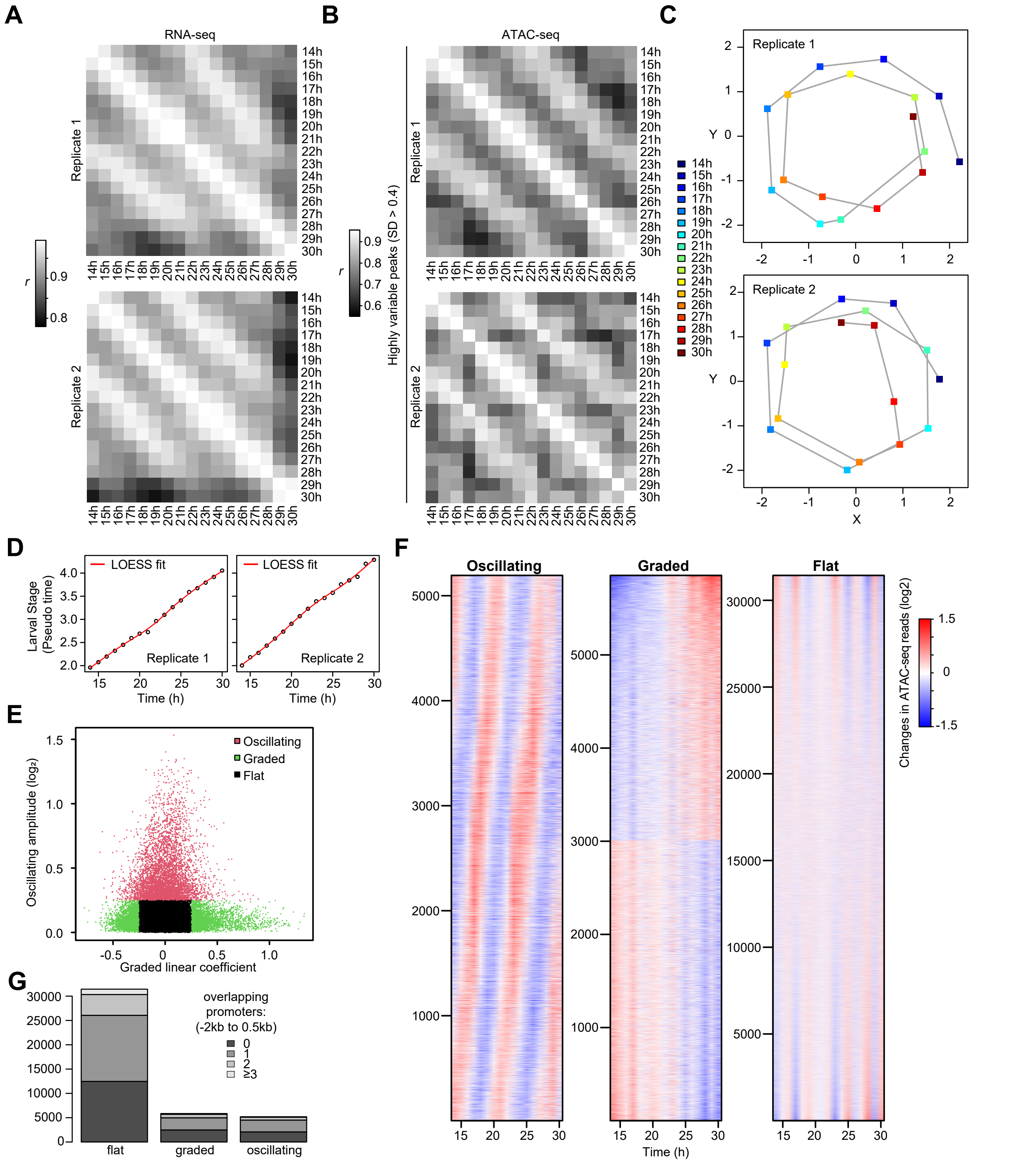


**Supplementary Figure 5**

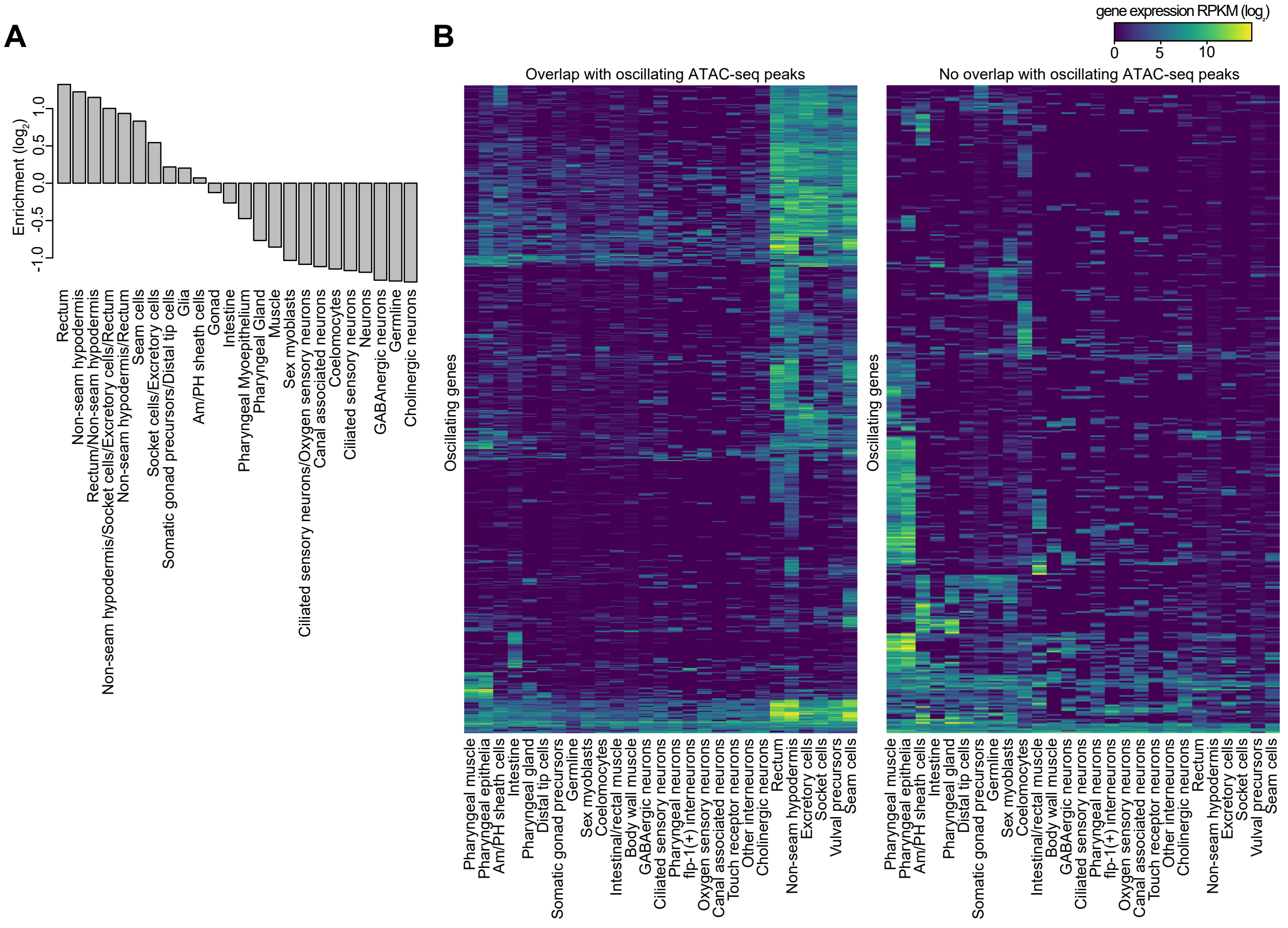


### Supplementary Figure 6

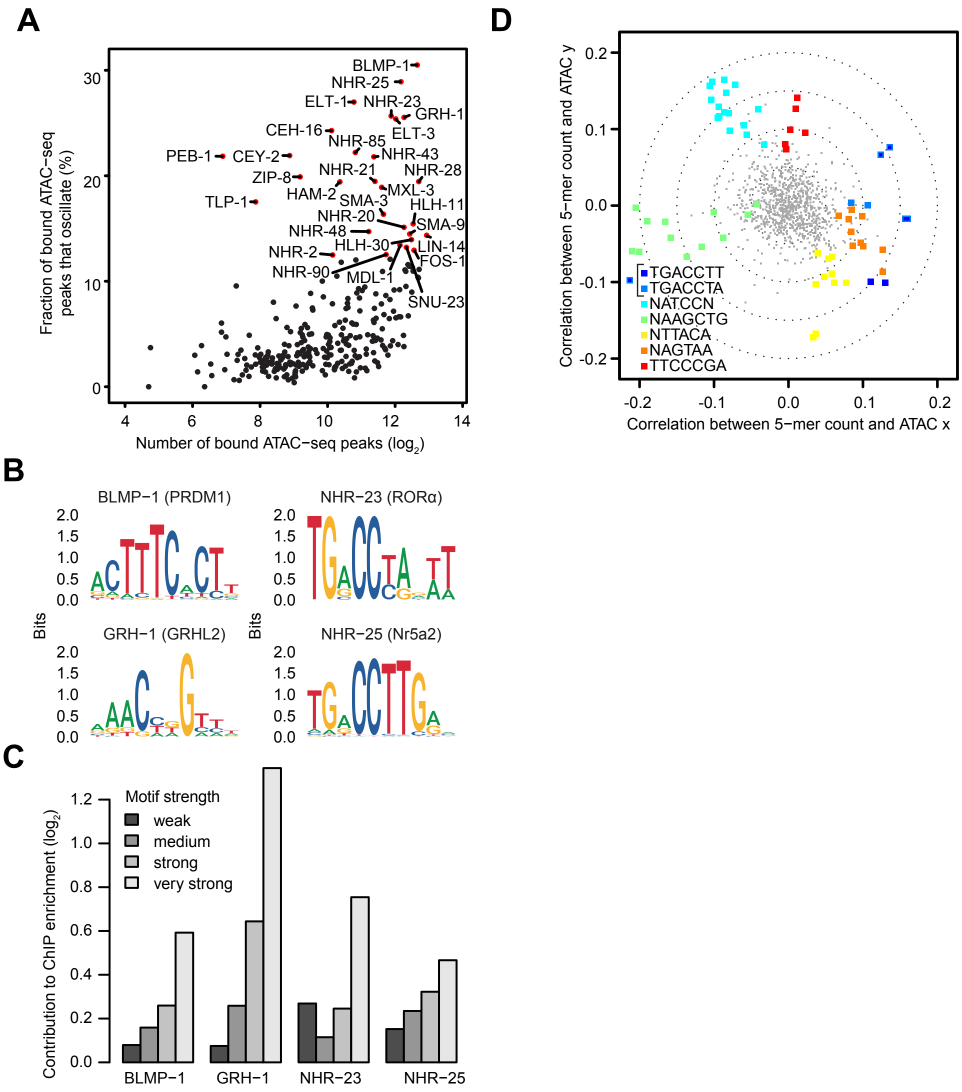


**Supplementary Figure 7**

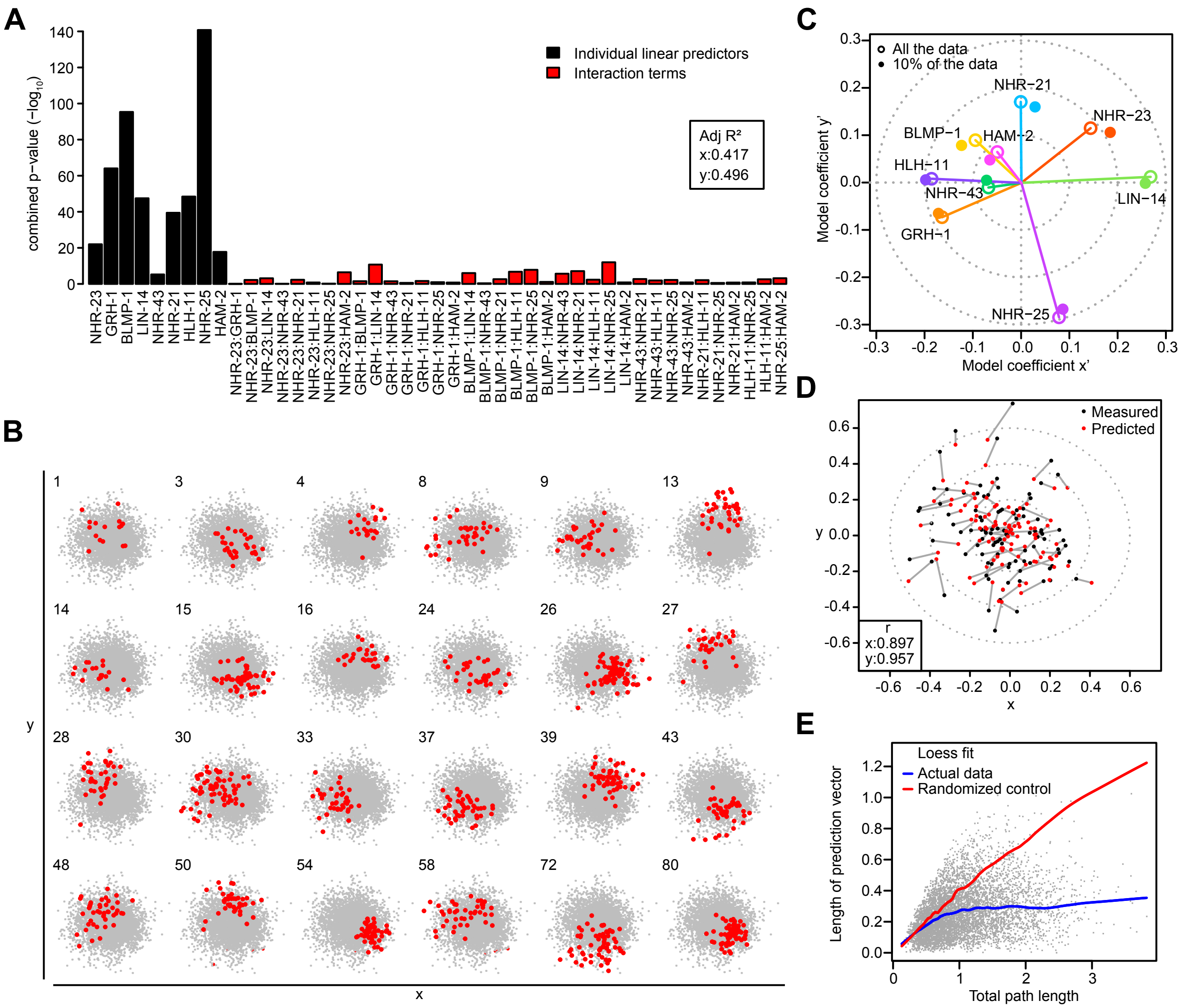


**Supplementary Figure 8**

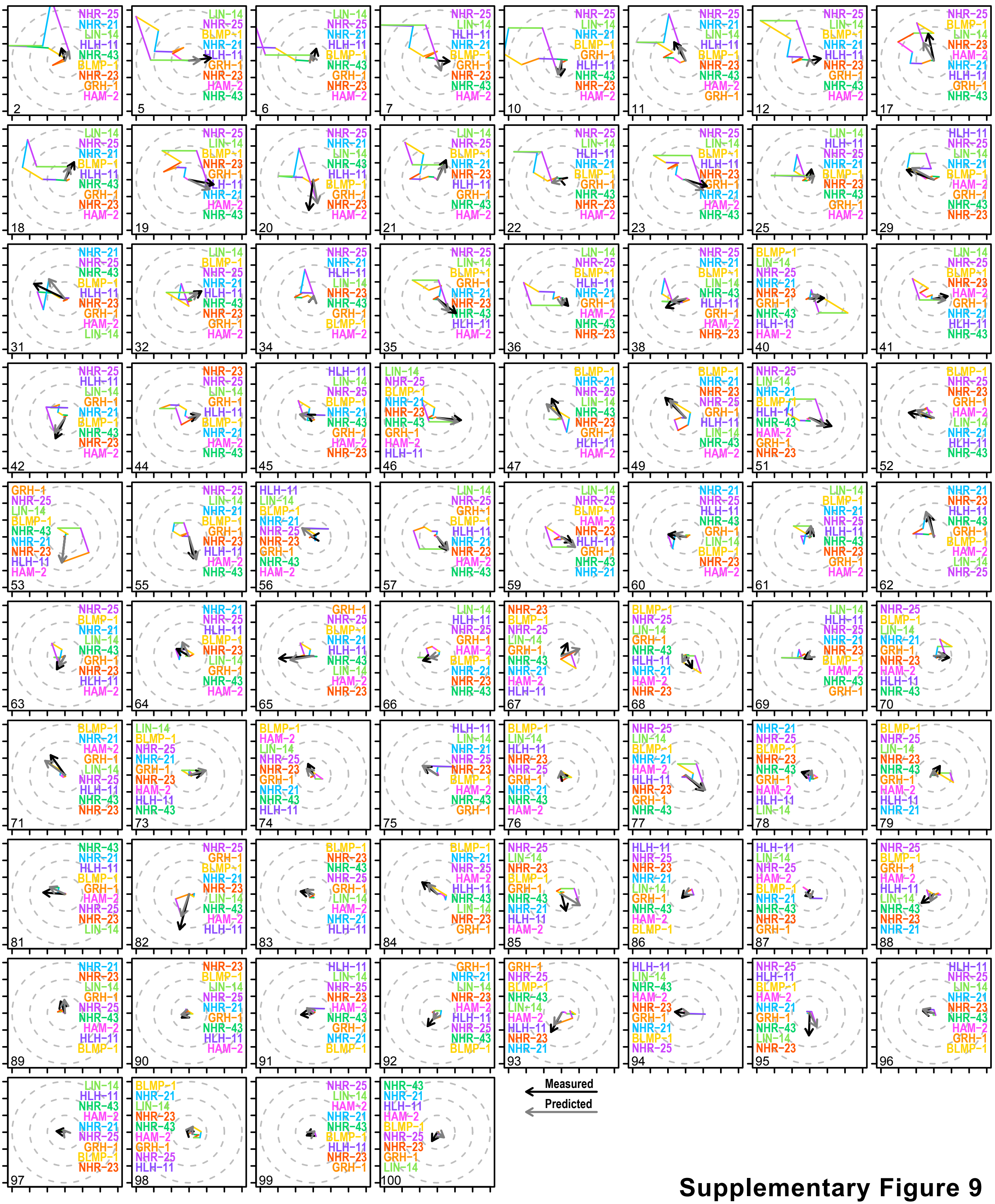


**Supplementary Figure 9**

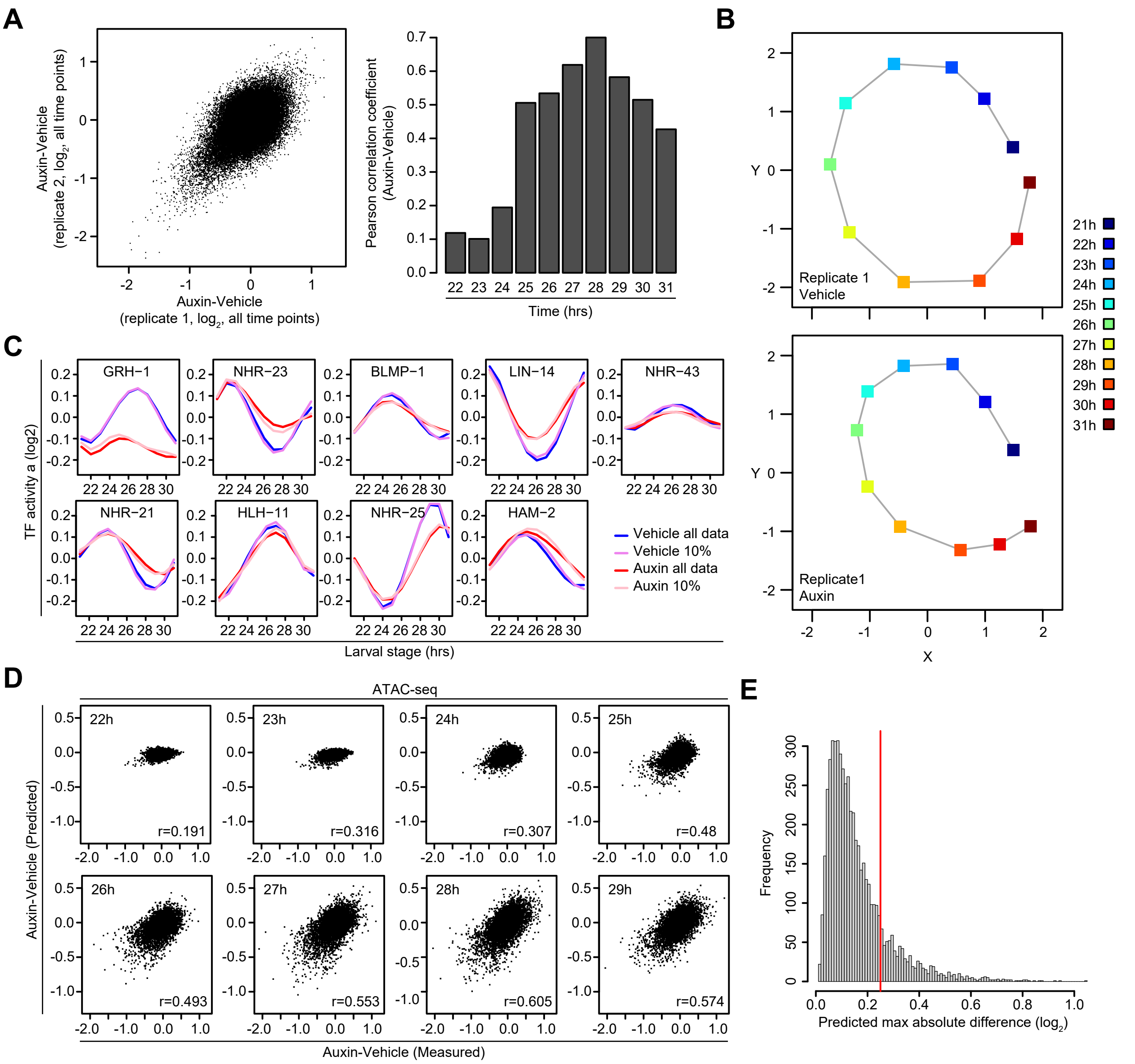


**Supplementary Figure 10**

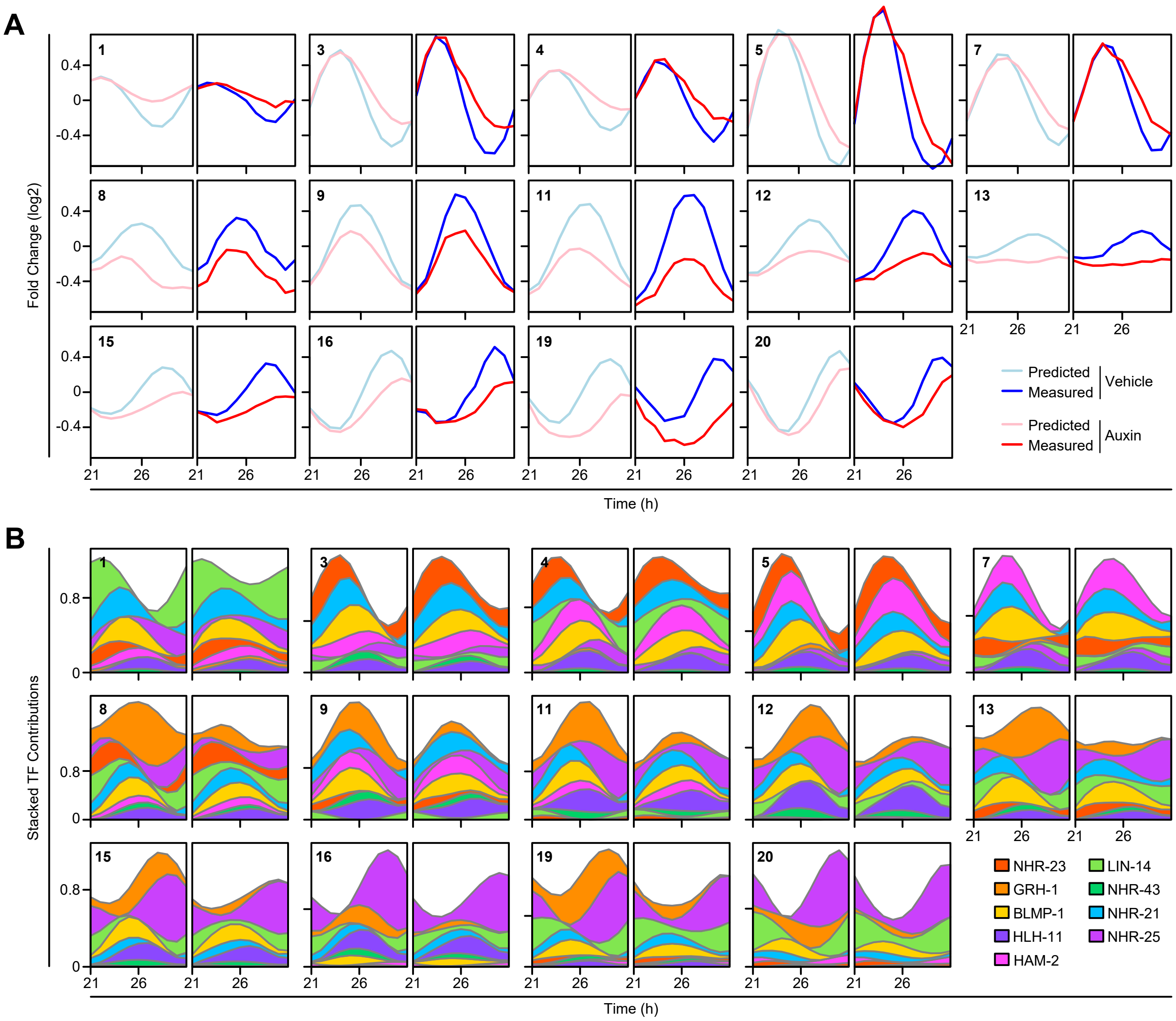


### Supplementary Figure 11
